# bigrig: A range simulator for the DEC[+J] model

**DOI:** 10.1101/2025.11.24.690345

**Authors:** Ben Bettisworth, Alexis Stamatakis

## Abstract

Quality software tools for science are, as a necessity, rigorously tested and verified. However, there is a major challenge to testing software used for phylogenetics and similar analysis. There is a shortage of ground truth data which can be used for validation, resulting in a reliance on simulated data. This reliance on simulated data results in a second challenge: the verification of the tool which generates the simulated data. In historical biogeography, software which simulated data has exclusively been implemented on an ad-hoc basis to verify specific tools. Here, we introduce `bigrig`, a simulator for the DEC[+J] model of range evolution. We show that `bigrig` is correct with extremely rigorous statistical testing and validation, ensuring that results deviate by no more than 0.0001 with 99.999% confidence. We also show that `bigrig` is extremely fast, capable of generating data for trees with tens of thousands of tips in under a second.

## 1 Introduction

Verifying the correctness of phylogenetics software is a perpetual challenge for tool developers (Darriba et al., 2018; Mendes et al., 2025; Carver et al., 2007; Bettisworth et al., 2023). The true phylogenetic tree—the series of speciation events originating from a common ancestor which explain an evolutionary history—is almost always unobservable. It is generally impossible to verify the results generated by software tools with a true, empirical reference phylogeny (but see Hillis et al. (1992) for one of the few notable exceptions.) As it currently stands, datasets for which the truth is known are exceptionally rare. Therefore, researchers cannot rely solely on these occasional known phylogenies to verify tool correctness.

A common goal in historical biogeography is to infer the ancestral ranges of species along a known phylogenetic tree (Varela et al., 2019; Baker and Couvreur, 2013; Vicente et al., 2017). However, similar to the true phylogenetic tree, these ancestral ranges generally can not be directly observed. Some ancestral ranges can be estimated from the fossil record (McLachlan and Clark, 2004), but there can be multiple complications which limit our knowledge about the true ancestral ranges. These complications include, but are not limited to: gaps in the fossil record; the study organism consists only of soft tissue which does not fossilize; or a geographical distribution which is unsuited for fossilization (Kidwell and Holland, 2002). Overall, there does not exist a reliable source of known ancestral ranges and associated phylogenetic trees.

Nonetheless, software verification remains vital, as without it is challenging to discriminate between a surprising result and an erroneous result which is the result of a software error. Therefore, one must turn to simulations in order to generate the necessary data to verify the correctness of the respective software tools (Mendes et al., 2025; Ly-Trong et al., 2022; Fletcher and Yang, 2009). Generally, artificial data are produced by assuming some statistical model of evolution, and subsequently generating datasets with a given set of model parameters. Data generated with this approach are unrealistic as they fail to capture the complexities of real biological systems (Trost et al., 2024). Despite this, the use of simulated data is still ubiquitous for software verification as currently no viable alternative exists.

However, software simulators are subject to the same concerns as inference software, specifically the presence of software errors can subtly corrupt the results. For example, these simulations might utilize the same code base for the inference of ancestral ranges and corresponding model parameters. In such cases, it becomes more likely that a software error in the model inference will also corrupt simulation results. Therefore, separating simulation code from inference code reduces the chances of integrated bugs in both the simulation and the inference. Additionally, it is critical that tools which implement inferences under a specific model are assessed using a common (simulation) standard. By using a common standard to evaluate tools one can ensure that improved performance (either computational performance or inference accuracy) is not merely a consequence of the evaluation process.

A common simulator also decreases the amount of work required to implement a new inference tool, as a fast and reliable method of generating testing data can be utilized by developers in the context of iterative development cycles. This will have the (eventual) result of increasing the number of tools available, and will increase the overall quality of tooling.

To date, biogeographical data simulations under the DEC[+J] model have often been implemented in an ad-hoc manner, with individual tools implementing their own simulation framework (Matzke, 2022; Bettisworth et al., 2023). This is done even though tools which can generate simulated data do exist (E.g. RevBayes (Höhna et al., 2016)). Indeed, RevBayes is capable of generating data under nearly any statistical model, but this comes at the cost of usability, as the user must specify the model themselves. This model specification task is non-trivial, and without expertise there is a high risk of inadvertently specifying an incorrect model. Therefore, instead of implementing ad hoc simulations for each software project, we can reduce the likelihood of errors by developing an independent stand-alone simulation tool.

In this paper we present `bigrig`, a simulator for the DEC[+J] model which complies with all of the aforementioned criteria for simulators. We explain the DEC[+J] model in detail in Section 2.1. Given a set of DEC[+J] parameters and a phylogenetic tree, `bigrig` will generate a range dataset for the taxa at the tips of the tree. In addition, ranges for inner nodes, cladogenesis events, and state transitions along branches will be generated. After generation, `bigrig` will log the results in a variety of practical file formats (YAML, JSON, or CSV), as well as output the results in phylip format, and an annotated tree in Newick format.

Furthermore, we show that `bigrig` is both highly reliable and extremely computationally efficient in regards to runtime and memory usage. We show that `bigrig` is reliable via two distinct approaches. First, we derive expected distributions for the results of fundamental model events, and perform statistical tests to ensure that the results are within 0.0001 with 99.999% of type 1 or type 2 error. Second, we produce an independent implementation which uses a slower, yet also simpler-to-implement method. Subsequently, we conduct statistical tests to verify that the results of these two independent implementations agree. Finally, we show that bigrig is computationally efficient. We demonstrate that `bigrig` can generate datasets for trees with tens of thousands of tips and 63 regions in under 0.5 seconds on a mid-range laptop.

## 2 Background

In the context of historical biogeography, an individual geographic sector where a taxon may be present is refered to as a *region*. A region can either be occupied, in which case we say it is full, or it can be unoccupied, in which case we say it is empty. A *range* describes whether each region is full or empty for a specific taxon for every region in a set. We denote the total number of regions for a range *r* as |*r*|. We write the number of full regions a range has as |*r*|_full_, and the number of empty regions as |*r*|_empty_. We denote the value of the *i*th region of a range as *r*_*i*_. In practice and for simplicity’s sake, we represent ranges as binary strings, with 1 indicating a full region, and 0 denoting an empty region.

For the regions *r* and *s* we denote the *bitwise and* operation as *r* ∧*s*, the *bitwise or* as *r ∨s*,the *bitwise exclusive or* as *r⊕s*, and the *bitwise negation* as *¬ r*. Here, *bitwise* indicates that the operation is executed independently on each character in the string (that is, each region). For example the *bitwise and* is defined as detail *r*_*i*_ *s*_*i*_ = *t*_*i*_ where 0 ≤ *i* ≤ |*r*|, 1 ∧1 = 1 and is equal to 0 otherwise.

These operations are of a particular interest as they can be efficiently computed via fundamental CPU instructions. Consequently, computing the results of these operations often requires less than a single CPU cycle^1^(Abel and Reineke, 2019). As such, computation of these operations (and functions which are composed of these operations) is extremely efficient.

### 2.1 An Overview of DEC[+J]

The Dispersion, Extinction, and Cladogenesis (DEC) model defines the probability of observing a given biogeographical history along a fixed phylogenetic tree with branch lengths. Under this model, there are three processes which govern range evolution. The first two, dispersion and extinction, model the stochastic range shifts over time, by modeling state changes that change a region to full or empty. The model assumes that each region may independently transition to a different state with waiting times which are exponentially distributed according to one of two rate parameters. For instance, with a full region *r*_*i*_ and extinction rate *e*, the waiting time *w* for *r*_*i*_ to transition to empty is *w* ∼ Exp(*e*). Alternatively, with a dispersion rate *d*, an empty region may transition to full with waiting time *w* ∼ Exp(*d*).

Cladogenesis, the third process in the DEC model, is when a parent species splits into two daughter species. During this event, full regions from the parent range are divided between the two daughters. The particular way in which parental ranges are inherited is restricted to a limited set of scenarios. In the original Ree et al. (2005) paper, these scenarios are simply given as Scenario 1, 2, and 3. The first two scenarios are intended to represent the pair of realistic cladogenic events, allopatry and sympatry. Scenario 3 covers the case when the parent range consists of a single full region, and so the two daughter species must inherit the same range. In the interest of clarity and memorability, we will use the names allopatry, sympatry, and copy to describe these scenarios.

Informally, the cladogenesis scenarios possible under the strict DEC model are: daughter ranges are disjoint (allopatry, alternatively vicariance); daughter species share at least one range (sympatry); daughter ranges are identical to the parent’s range, and all regions are singletons (copy). We provide a formal definition of these cladogenesis events in Section 2.2. Matzke (2014) extended the set of cladogenesis events by incluidng a novel “jump” scenario that aims to represent a “founder-event speciation” event. Here, a small population becomes isolated by colonizing a novel region. Additionally, Matzke (2014) also allowed for the relative probability of the cladogenesis events to be inferred parameters, such that they may vary between datasets. Under this extension, sympatric events might have a larger weight, and therefore exhibit a higher likelihood when observed, than allopatric events. These extensions are generally denoted by DEC+J, similar to how gamma rate categories are denoted by GTR+G4 in standard phylogenetic models.

The parameters of the strict DEC model are the dispersion and extinction rates (*d* and *e*), and a fixed phylogenetic tree with fixed branch lengths. DEC[+J] adds the cladogenesis parameters *s, v, y* and *j*, for sympatric, allopatric (vicariance), copy, and jump events, respectively.

### 2.2 A Formal Definition of Cladogenesis Events

We will represent a cladogenesis event as the tuple (*p, l, r*), where *p* is the parent range, and *l* and *r* are the left and right child ranges, respectively. The DEC[+J] model assumes that specitation (and therefore range inheritance) occurs at an instant in time and is restricted to a single region. This implies that at least one of the daughter ranges (*l* or *r*) will be a singleton. For the remainder of this section and without loss of generality, we assume that *l* is the singleton range.

When a cladogenesis event occurs, the *strict* DEC model assumes equal probability for all event types. In other words, when considering if a particular cladogenesis event is more likely to be allopatric or sympatric, the *strict* DEC models these as being equally likely. To compute the probability of a type of cladogenesis event over the set *T* =*{* Allopatry, Sympatry, Copy *}* we compute

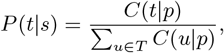

where *C*(*t* | *p*) is the count of possible cladogenesis events of type *t* given *p*, and *T* is the set of *all* possible event types.

This cladogenesis model is extended in DEC[+J] by including weights and an additional cladogenesis type. Under this extended model, we compute the probability as

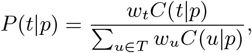

where *w*_*t*_ is the weight of cladogenesis type *t*. Additionally, *T* is augmented by the new type “jump” to become *T* =*{*Allopatry, Sympatry, Copy, Jump*}*. Informally, “jump” events are cladogenesis events which allow one of the daughter species to disperse to an unoccupied range.

We define the cladogenesis event type as

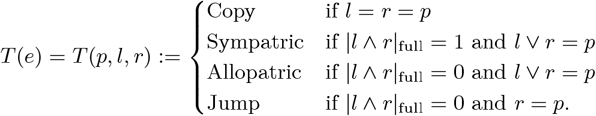

## 3 Methods

For generating data under the DEC[+J] model, two simulation phases are required. Range changes occurring along a branch (dispersion and extinction) and ranges that occur as a result of speciation events (cladogensis) need to be simulated as separate processes. For mnemonics, we will refer to these phases as the *spread* and *split* phases.

As already mentioned, for both phases, we implemented two completely independent data simulation methods in order to verify that the results of these two distinct implementations are numerically identical. The first simulation method (for both the spread and split phases) is based on the design pattern of rejection sampling. In rejection sampling, samples of a distribution over a complicated sample (say Ω) space are generated by initially sampling from a distribution over a simpler sample space (say Ω^*I*^). If a sample drawn from Ω^*I*^ is not in Ω, then the sample is rejected as invalid, and we draw a new sample. This process is repeated until we draw a valid sample^2^. Rejection sampling can be modified to draw non-uniform distributions by means of rejecting valid samples by some specified proportion in order to correct for the bias. The precise rejection probability in this case depends on the specific distributions. We will specify the rejection probability in detail when we discuss each method.

Rejection methods are extremely simple, and therefore easy to implement correctly. The unfortunate trade-off is that a substantial amount of time is spent on generating ultimately rejected samples. The time spent generating invalid samples is ultimately wasted. Therefore implementing software will necessarily be, to some degree, computationally inefficient. Rejection sampling is typically only utilized when no other method is capable of producing acceptable results.

In addition to the rejection methods, we implemented “fast” methods of generating samples for the spread and split phases. These “fast” methods implement two major improvements: analytic and CPU-aware optimizations. Here, an analytic optimization relies on a fast algorithm which will analytically generate samples from the desired distribution. However, we need to keep in mind that the analytical correctness of an algorithm does not imply that it is *numerically* correct (Goldberg, 1991; IEEE, 1985). In addition, we implement CPU-aware optimizations which leverage the computational capabilities of modern CPUs.

We can then use these independent implementations (rejection and “fast” method) to verify the simulation results. By verifying that the results of the two methods are equivalent, in a rigorous statistical sense, we can deduce that if there is a software bug, it must be present in both methods. However, the probability of the same software bug being present in two independent and algorithmically distinct different implementations is extremely low (Sklaroff, 1976; Taneja et al., 2010). Therefore, if the results of the two methods produce statistically equivalent results, we can be confident that the results are correct.

### 3.1 Simulating the Spread

In the DEC[+J] model, the dispersion and extinction processes are modeled as a continous time markov chain (CMTC). CMTCs are themselves a generalization of many independent Poisson processes. We use this fact to draw samples to generate valid spread events. A spread event *e* is defined as the tuple (*p, c, w*), where *p* is the parent range, *c* is the child range, and *w* is the waiting time between *p* transition to *c*. For any spread event *p* and *c* differ by in only one region. More formally |*p⊕ c*|_full_ = 1.

The problem is this: given a range *r* with |*r*| = *n* regions and branch length *t*, produce a final range which has undergone the processes of extinction and dispersion at rates of *e* and *d*, respectively. Each region experiences either extinction or dispersion independently of each other. This observation is the basis for the rejection method of spread simulation: sample *n* waiting times, labeled *w*_*i*_, one for each region. Specifically, the waiting times are distributed as

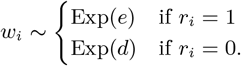

Once all *n* waiting times have been sampled, we find *i* = argmin(*w*_*i*_) and negate the corresponding region so that 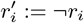. The resulting transition event is (*r, ŕ, w*_*i*_).

We repeat the above process until the total waiting, the sum of selected *w*_*i*_’s, time exceeds *t*, at which point the process halts, and we yield the list of transition events, excluding the final generated event.

Keen readers might already be aware of the fact that if we have a set of independent processes as above, the entire set can be represented via a single exponential distribution, with *w* ∼ Exp(*q*) where

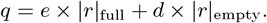

This second method then is to compute *t* using Eq. 3.1, draw a waiting waiting time *w* ∼ Exp(*t*). Once a time is sampled, we pick a region with weights equal to the exponential distribution parameter in Eq. 3.1.

We implement both of the above methods, but only use the second to generate samples in by default. The first method, the rejection method, is used to check the results of the second method, the faster method.

### 3.2 Simulating the Split

A split will be written as an ordered triplet of ranges (*p, l, r*), where *p* is the parent range and *l* and *r* are the respective child ranges. Given *p*, we can sample a split by sampling two random numbers from a uniform distribution, and rejecting invalid splits. That is, we accept splits which are sympatric with the probability

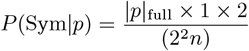

as there is one way to pick a region which equals *p*, and there are |*p*| _full_ ways to pick a region which is both a singleton and a subset of *p*. Furthermore, the left and right child can be swapped.

By similar logic, there are *p* _full_ ways to pick a region such that |*l* |_full_ = |*p*| _full_*−* 1. Once *l* has been picked, there is only one region which will produce a valid split event. Therefore,

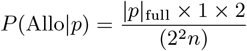

which is equal to *P* (Sym *p*). Therefore, for distributions where *j* = 0 and *s* = *v*, the probability of drawing either a sympatric or allopatric event is the same. Additionally, since the probability of any particular *l* and *r* is equal given a split type, then all valid splits have the same probability.

In the case when *j* ≠ 0, then

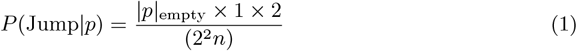

In order to implement the weighted split type introduced by Matzke in Matzke (2014), we accept a sampled split event with probability that is equal to the normalized split weight. For example, if the split type weights are *y* = *s* = *v* = *j* = 1.0, then a jump type split will be accepted with probability *j/*(*y* + *s* + *v* + *j*).

To accelerate this process, we implement an optimized split simulation procedure. First, we generate the split type according to the relative split event weights (*y, s, v*, and *j*). Sampling a split is divided into two cases, the singleton case where |*p*| _full_ = 1, and the non-singleton case where| *p*| _full_ *>* 1. In the singleton case, we generate a split type from *{*jump, copy*}*, weighted accordingly. In the non-singleton case, we generate a split type from *{*jump, sympatry, allopatry*}*, also weighted accordingly.

We compute the weight for each type as

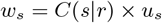

where *C*(*s*| *r*) is the count of feasible splits of type *s* given a region *r*. For example, for region 11 there exists two ways to split allopatrically: (10, 01) and (01, 10) The exact formulas for various types are

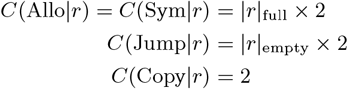

Once a split type has been sampled, we generate a the daughter ranges according to the type by randomly picking which region will be flipped. If the split type is a jump, the region is chosen from among the empty regions. If the split type is either allopatry or sympatry, the region is chosen from the full regions. Once the region is flipped and two regions are produced, we randomly determine which child obtains the singleton region, with probability 0.5.

#### 3.2.1 Simulating a Phylogenetic Tree

Given a root range *r*, and parameters *d, e* and *c*, and an age *t*, we can sample a phylogenetic tree with Algorithm 1. In short, a tree is sampled by starting with a root range *r* and a tree age *t*. The range *r* is then evolved under the DEC[+J] process. Waiting times until an event, either a spread or split event, are sampled, and then the corresponding event is generated with *r* as the parent range. When the waiting time for a branch exceeds *t*, the process halts. We recursively iterate over all branches until the path from each extant tip to the root has a length of *t*.

##### Algorithm 1

Sample a tree under DEC[+J], given a deadline *t*.

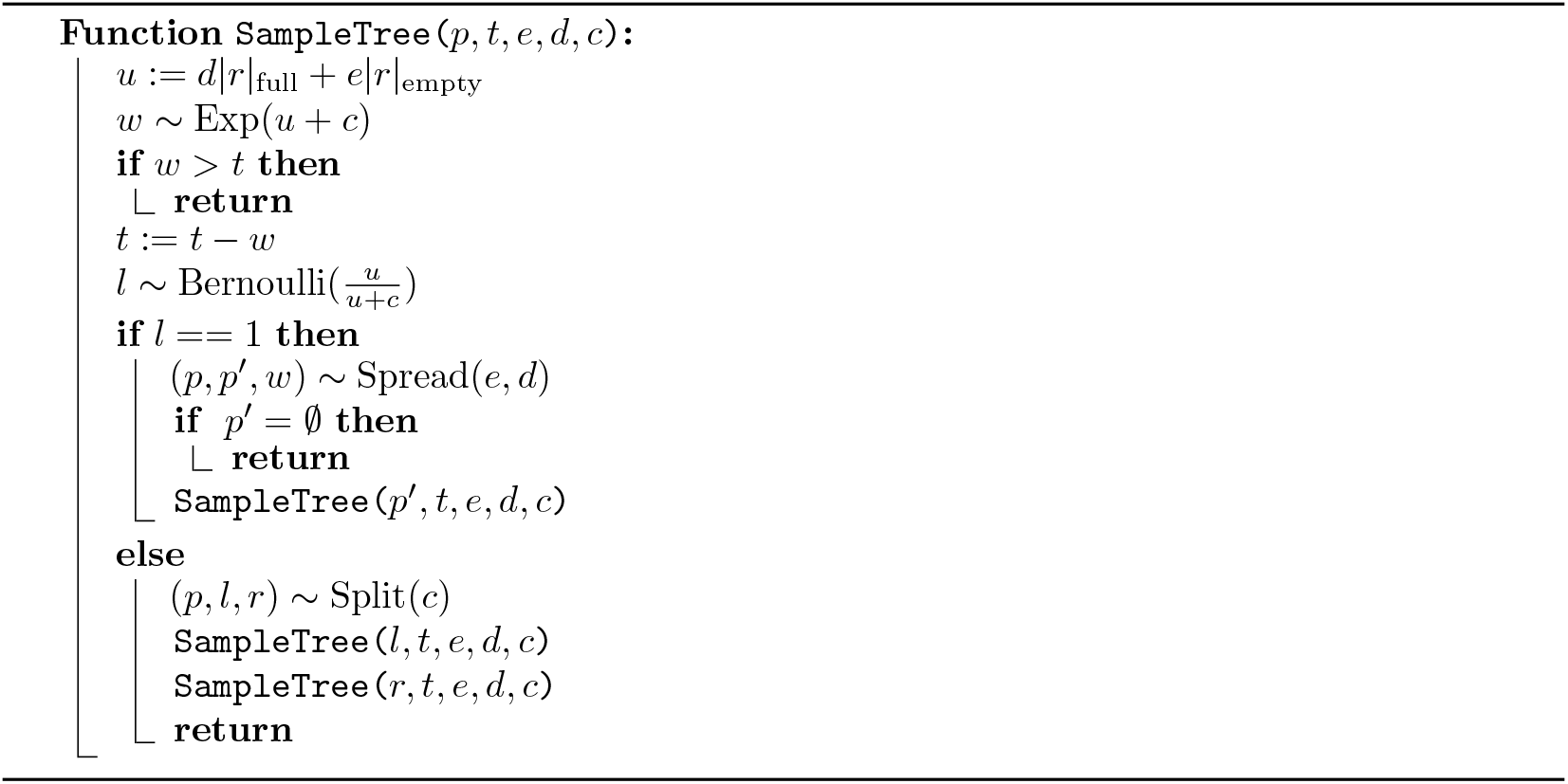

### 3.3 Testing

Our primary strategy for testing for correctness is to test the two processes, split and spread, independently with statistical rejection testing with extremely large sample sizes to minimize the chance and degree of error. To this end, we implemented our tests in C++, using the core functions of `bigrig` as a library. Writing the tests in a performant language like C++ allows us to compute large sample sizes in a reasonable amount of time.

#### 3.3.1 Spread

For ensuring correct results and implementations, we conduct two types of tests. The first test computes the expected value of the waiting time given the model parameters analytically, and performs a t-test against that expected value. We compute 1,886,084,219 samples of the spread distribution, which allows us to be 99.999% confident that the error is less than 0.0001. We perform this test for 7 different initial ranges with, each with a size of 4 regions, and 16 different parameter sets, for a total of 116 separate tests.

The second test is a regression test against the rejection method. We again run both methods for 1,886,084,219 iterations, which again ensures that the error is less than 0.0001 with 99.999% confidence. Furthermore, we also perform this test for 5 different initial ranges, each with a size of 4 regions, and 16 different parameter sets, for a total of 80 separate tests. We perform fewer tests due to the increased computational cost of the rejection method.

Finally, we conducted a *χ*^2^ test to ensure that regions are being picked at the expected rate. For this test, we enabled either dispersion or extinction, including both, and computed the expected number of substitutions for each region given the initial range. With this initial region, we sampled 1000 spread events, and the utilized a *χ*^2^ test to ensure the realized distribution matches the expected distribution. We reject the hypothesis that the two distributions are equivalent with *α* = 0.0001, I.E. we reject with 99.99% confidence.

#### 3.3.2 Split

We compute a sample of 2,019,696,124 split events, and compare this sample against the expected distribution of cladogenesis event types using a G test. This gives a probability of either type I or type II error occurring at 0.00001, with a maximum deviation from the expected value of 0.0001. Please see the supplemental material for full derivation of these values. We perform a test for each of 6 initial ranges and 5 cladogenesis parameters for a total of 30 tests.

Likewise, we also compare the results of the rejection method with the analytical method. Due to the expensive nature of rejection sampling, we limit our sample size to 201,970 split events from each of the methods. We then use the same G test to compare the results. We perform this test for 6 initial ranges and 5 sets of cladogenesis parameters for a total of 30 tests.

Finally, we conducted a *χ*^2^ test to ensure that regions are distributed as expected between the child regions. For event types Sympatry, Allopatry, and Jump, we sample a split event of the specified type and determine the singleton region. We then ensure that realized counts match the expected counts with a *χ*^2^ test with rejection threshold of *α* = 0.0001.

### 3.4 Benchmarks

We also sought to evaluate the performance of `bigrig`. In our initial investigations, we found that the time to generate data was dependent on several model parameters, in particular the tree height and the cladogenesis parameters. Therefore, we sought to assess both the typical performance and the extreme performance of `bigrig`. We generated 10 random rooted ultrametric trees with 2^*i*^, 3≤ *i*≤ 16 each, for a total of 140 trees. These trees were generated via a Yule process, with the branch lengths scaled so that the tree height was equal to 1.0. For each tree generated, we generated 10 random ranges of size 3, 7, 15, 31, along with the CMTC model parameters *e, d*∼ Uniform(0, 1) and cladogenesis parameters from either without jumps (*s* = *v* = *y* = 1, *j* = 0) or with jumps (*s* = *c* = *y* = *j* = 1). The first cladogenesis parameter set is the fixed parameters used in the original DEC, whereas the second parameter set simply “enables” jumps to occur, I.E. converts the model to DEC[+J]. This yields 10 datasets for each tree, 20 datasets for each range size, for a total of 14,000 datasets. We implemented this pipeline in Snakemake(Mölder et al., 2021) and we measured execution times with the Linux tool `perf`. The execution times of `bigrig` tend to be very short, so to measure with sufficient resolution we found it was required to utilize perf which uses kernel and hardware level performance counters to measure runtime.

In addition to the performance benchmarks, we also benchmarked the software quality using Softwipe v0.2 (Zapletal et al., 2021). Softwipe assess the code quality via a series of hureistics, and then produces a score between 0 and 10, with 0 being low quality, and 10 being high quality.

## 4 Results

For all tests we conducted, `bigrig` passed. That is, all methods produced results which were equivalent between the rejection and analytic methods, and each analytical methods produdo we have a list of softwipe ced the expected distribution. As each statistical test can be considered independent, we can estimate the probability of an implementation error in `bigrig` going undetected as the product of a type I error for each test. To be explicit, the probability of an undetected error is nearly 0, and we can be nearly certain that no such bug exists in `bigrig`.

In general, `bigrig` is extremely fast, as can be seen in the Fig. 1. The mean time for execution varies from 0.030 seconds (*SD* = 0.002) with 3 regions and 65536 taxa to 0.046 seconds (*SD* = 0.004) with 63 regions and 65536 taxa. The minimum time over all trials was 0.000004 (3 regions, 8 taxa) seconds, and the max time was 0.081 seconds (63 regions, 65536 taxa). The overall execution times are shown in Fig. 1.

**Figure 1:**
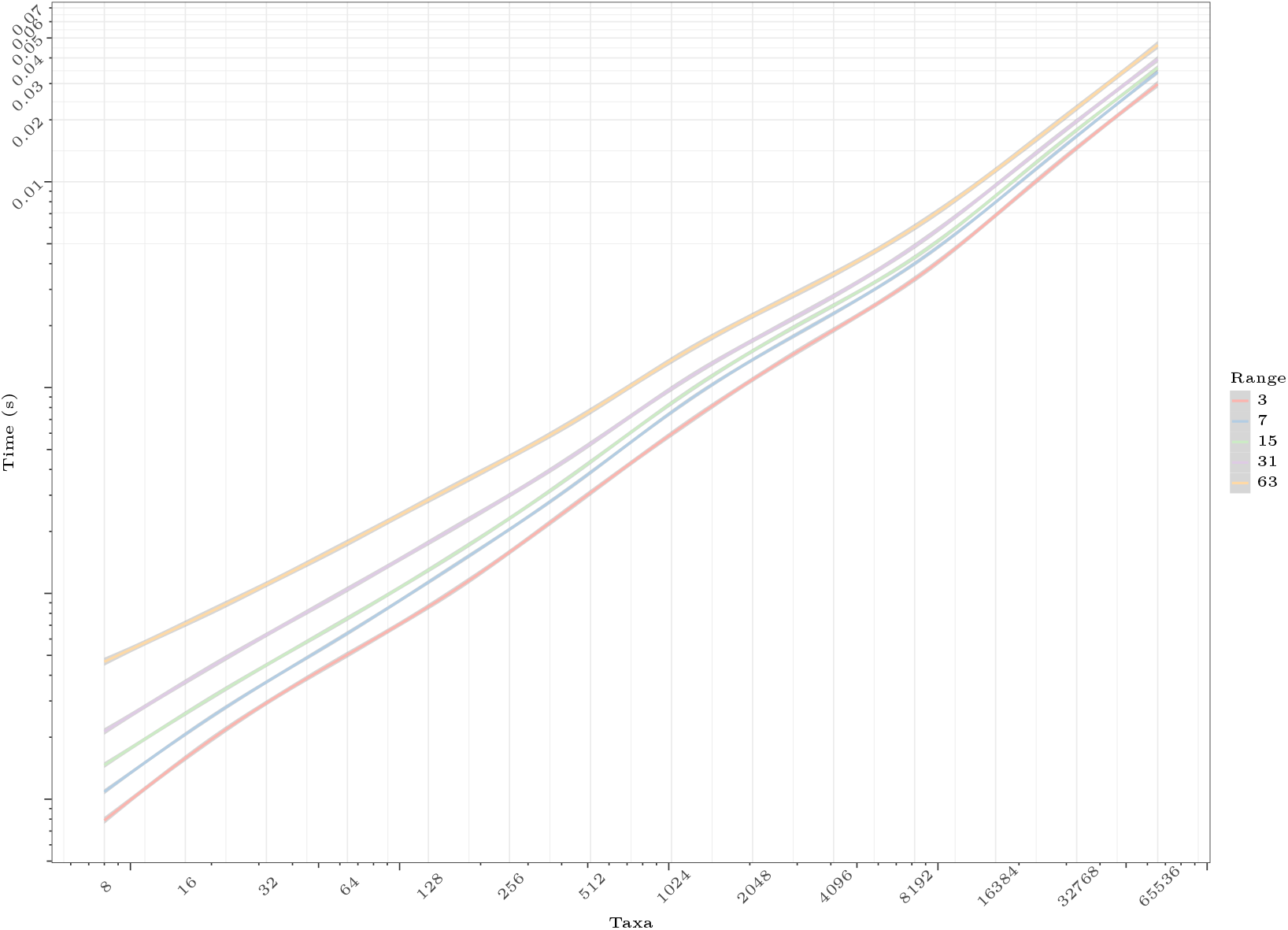
Plot of smoothed generalized additive model (GAM) fit to runtime data for bigrig execution on a log-log scale. Plot was generated with `ggplot2’s geom_smooth()` function. For each taxa quantity, 10 random rooted trees were generated via a Yule process with branch lengths scaled to a total tree hight of 1.0. For each tree, 10 sets of CMTC model parameters were randomly sampled such that *e, d* Uniform(0, 1). Cladogenesis parameters were set either to *s* = *v* = *y* = 1.0 and *j* = 0.0 or *s* = *v* = *y* = 1.0 (see text for details). Shaded area indicates the 0.95 confidence band as computed by the `predict.Gam` function in R.

The jump parameter had a noticeable, but small, impact on the overall runtime of `bigrig`. The mean time to generate data with 63 regions and 65536 taxa when *j* = 1 was 0.048 (*SD* = 0.002) versus 0.044 (*SD* = 0.005) with *j* = 0. This can be seen as well in Fig. 1, as the increase in execution time from added regions or added taxa is much larger than the standard deviation (the grey shaded region). Additionally, the difference between jumps and no jumps is statistically measurable via a two sided t-test *p≈*8*×*10^*−*12^.

Softwipe evaluated the `bigrig` code to have a software score of 8.5, indicating a very high quality of software. This indicates that `bigrig` is among the tools with the highest software quality.

## 5 Conclusion

We have presented `bigrig`, a reliable, fast, and easy to use tool to generate simulated historical biogeography datasets. We have shown that `bigrig` has a negligible chance of having a software implementation error. We have also show that it is extremely fast, capable of generating a dataset in under a second which is far beyond the ability of any current inference software to infer.

In the future, we wish extend the features of `bigrig` to include dataset generation for the family of State Specific Evolution (SSE) models (BiSSE, GeoSSE, ClaSSE) (Cornuault and Sanmartín, 2022). The primary purpose of these extensions is to test the robustness of DEC[+J] models to model violations, in order to asses the added value of these models. This analysis is important for future tool development, as the more advanced SSE member models cannot be inferred using the typical CMTC computational framework use for DEC[+J]. However, at this time it seems like these models will exhibit similar scaling during inference as DEC[+J] as all models here share the exponential scaling of their state space.

Investigating exactly when DEC[+J] fails to give adequate results will help both researchers and tool developers to efficiently utilize their scarce resources. In the case of users, it will help conserve compute time, and in the case of tool developers it will help conserve developer time.

## 6 Acknowledgements

Many thanks to Alexander I. Jordan from the Computational Statistics group at the Heidelberg Institute for Theoretical Studies for his assistance on deriving and implementing the statistical tests used to verify the results of `bigrig`.

This work was funded by the European Union (EU) under Grant Agreement No. 101087081 (Comp-Bio-div-GR).

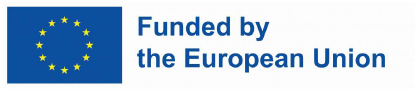

## 7 Data Availability

The releases and code for `bigrig` are available at github.com/computations/bigrig. The scripts used to benchmark `bigrig` and plot the results are available at github.com/computations/bigrig-test.

## Footnotes

1 This is due to the capabilities of modern CPUs to potentially execute more than one instruction per cycle, depending on the which instructions are executed.

2 This process can fail to terminate with probability 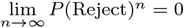

